# A novel small molecule PKC epsilon inhibitor reduces hyperalgesia induced by paclitaxel or opioid withdrawal

**DOI:** 10.1101/2023.06.01.543325

**Authors:** Adriana Gregory-Flores, Ivan J. Magayewski Bonet, Stève Desaivre, Jon D. Levine, Stanton F. McHardy, Harmannus de Kraker, Nicholas Russell, Caleb Fleischer, Robert O. Messing, Michela Marinelli

**Author notes:** Equal contribution (Robert O. Messing, Michela Marinelli). Corresponding Author: Michela Marinelli, Ph.D.; Mailing address: 1601B Trinity Street, HDB 5.322, Austin, TX 78712.

## Abstract

Aggressive marketing and increased prescribing of opioids for treating pain have fueled a prominent increase in opioid use disorder. This acute public health problem has led to calls for the development of non-opioid alternatives to treat chronic pain. The enzyme protein kinase C epsilon (PKCε) plays an important role in nociceptor sensitization, in inflammatory and neuropathic pain. Here we investigated the effects of a novel small molecule that inhibits PKCε in a rodent model of chemotherapy-induced neuropathic pain produced by administration of the cancer chemotherapy, paclitaxel. Because transition of opioid-dependent individuals to a non-opioid pain medication can increase pain due to opioid withdrawal, we also investigated if the PKCε inhibitor alters features of opioid withdrawal and opioid self-administration. This novel PKCε inhibitor attenuated paclitaxel-induced hyperalgesia, reversed hyperalgesia produced by opioid withdrawal, and reduced somatic signs of opioid withdrawal. This PKCε inhibitor did not modify opioid self-administration, nor produce self-administration. These findings suggest that PKCε inhibition is an effective, non-addictive strategy to treat chemotherapy-induced neuropathic pain with the added benefit of limiting hyperalgesia due to opioid withdrawal, which could facilitate switching treatment of chronic pain, in opioid-dependent individuals.

## Introduction

According to the Centers for Disease Control and Prevention (CDC), an estimated 50 million adults in the United States experience chronic pain (1), with an estimated annual cost exceeding $600 billion (2, 3). The quality of life for these individuals is impaired due to significant suffering, disability, and unemployment.

For patients with chronic pain, pain management emphasizes psychological treatment, physical therapy, lifestyle modification and education in self-care, with recommendations for analgesia that include NSAIDs, antidepressants, and anticonvulsants; opioids are reserved for severe or disabling pain that is insufficiently controlled by non-opioid drugs (4, 5). Unfortunately, these medications have serious side effects that limit their chronic use (6). With chronic use, NSAIDs can cause gastrointestinal bleeding and renal insufficiency, and those that are selective inhibitors of COX-2 carry increased risk of cardiovascular events. Antidepressants can cause autonomic dysfunction and, in high doses, can produce arrhythmias, hypotension, and seizures. The anticonvulsants gabapentin and pregabalin produce somnolence, fatigue, dizziness, ataxia, and weight gain. And opioids are sedating, and frequently cause nausea and constipation. They also induce tolerance, requiring increases in dosage, and in high doses, they produce respiratory depression, which can be fatal. Furthermore, discontinuation of opioids in people taking them chronically for pain management produces aversive withdrawal signs and hyperalgesia, further complicating pain management. Finally, repeated opioid use can lead to addiction in 8-12% of patients (6). Because of these limitations and a lack of efficacy, particularly for the treatment of chronic pain, there is still a major need for developing more effective pain medications with improved side effect profiles.

A particularly difficult to treat class of pain syndromes are those resulting as a common side effect of cancer treatment: chemotherapy-induced peripheral neuropathy (CIPN) (7, 8). CIPN adversely affects the quality of life in oncology patients due to discomfort, pain, disability, and risk of falling. With cytotoxic agents such as the chemotherapeutic agent paclitaxel, pain is common and can be persistent with approximately half of patients having symptoms one year after treatment (7, 9). Unfortunately, CIPN-induced pain is poorly responsive to available medications and the only agent recommended by the American Society of Clinical Oncology is duloxetine, which has modest effects (7, 10).

Protein kinase C is a family of nine serine-threonine kinases that transduce signals carried by lipid second messengers (11). A large body of literature indicates that one PKC isoform, PKCε, is particularly important for signaling in peripheral nociceptive sensory neurons, and that this kinase mediates inflammatory, neuropathic, and stress-related pain, and the transition from acute to chronic pain (4, 12–14). Such findings have sparked interest in developing small molecule inhibitors of PKCε to treat chronic pain.

Here we report the discovery of a small molecule with drug-like properties that potently inhibits PKCε and reduces paclitaxel-induced hyperalgesia in rats. Surprisingly, this compound also prevented and reversed hyperalgesia provoked by opioid withdrawal. The compound was not self-administered and did not alter opioid self-administration. Our results identify a small molecule targeting PKCε that has potential to become a new class of non-addictive analgesics to treat CIPN and hyperalgesia evoked by opioid withdrawal.

## Results

### Characterization of the PKCε inhibitor CP612

We previously reported the design of PKCε inhibitors (15) based on Compound 397 (described in patent WO 2007/006546), which was developed from the ROCK inhibitor Y-27632. The lead compound we tested (Compound 1) inhibited PKCε but enhanced the hypnotic effect of ethanol in *Prkce* null mice, indicating an off-target effect. This compound also transiently depressed locomotion (15) in association with reversible hypotension (data not shown). We suspected these effects were due to ROCK inhibition since ROCK inhibitors can cause hypotension (16, 17) and Compound 1 inhibited ROCK1 (IC50 = 318 ± 40 nM; n = 38) and PKCε (IC50 = 120 ± 16 nM; n = 38) with similar potency.

To improve selectivity and potency for PKCε, we carried out structure-activity-relationship studies that led to the development of CIDD-0150612 (CP612), which possessed known favorable physicochemical properties for central nervous system (CNS) drugs (18). This compound inhibited PKCε with IC50 = 1.1 ± 0.7 nM (n = 35) and ROCK1 with IC50 = 43.7 ± 4.2 nM (n = 36) making it ∼ 40-fold more selective against PKCε over ROCK1. CP612 was screened for kinome specificity at 200 nM against 468 kinases using the scanMax assay panel from Eurofins-DiscoverX. It inhibited 10 other wild type kinases besides PKCε, all to < 10% of control: CLK1 (CDC Like Kinase 1), CLK4, PKN1 (Protein kinase N1), PKN2, PKCδ, PKCη, PKCθ, PRKG2 (Protein Kinase CGMP-Dependent 2), ROCK1, and ROCK2. When assayed against several PKC isozymes, CP612 did not inhibit atypical PKCζ, weakly inhibited conventional PKCβII and PKCγ, and within the nPKC subfamily, was most potent against PKCε (Figure 1A).

**Figure 1.**
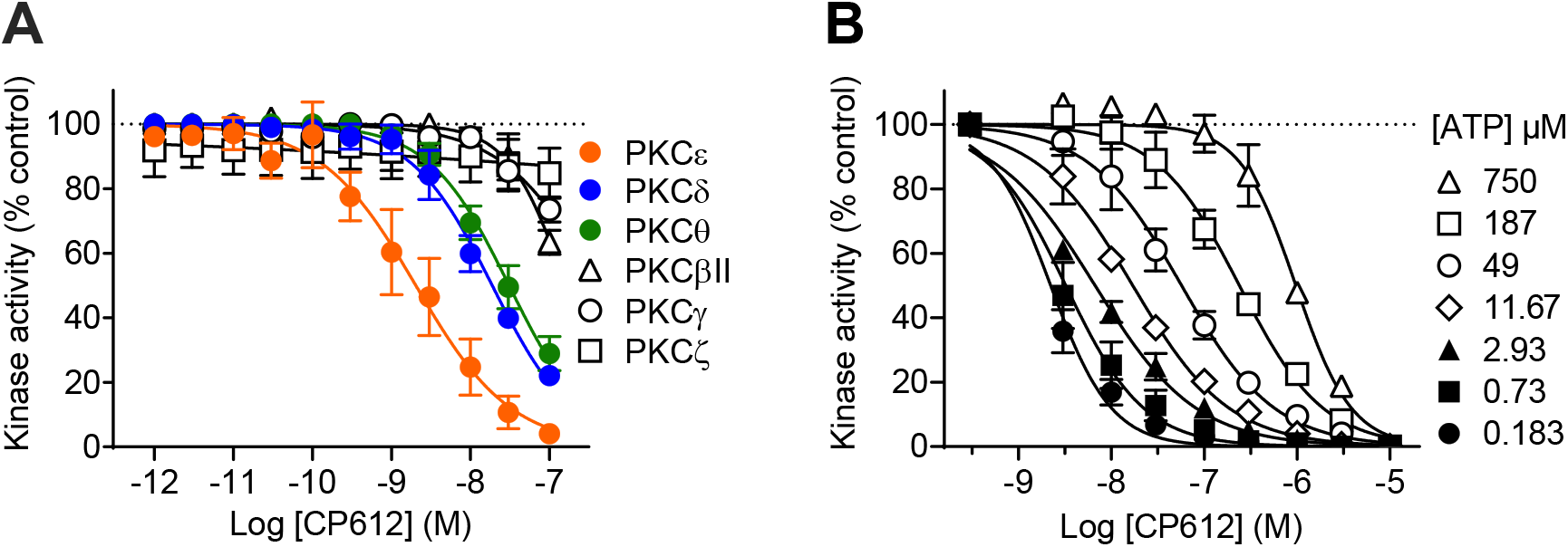
Characterization of CP612. **(A)** CP612 inhibited novel PKCs by > 50% at 30 nM but was more potent against PKCε (IC50 = 2.03 nM) than PKCδ (IC50 = 18.8 nM) or PKCθ (30.7 nM). **(B)** Dose-response curves for inhibition of PKCε by CP612 showing a rightward shift at higher [ATP]. Data are mean and SEM (n = 3 experiments each done in duplicate or triplicate).

Since CP612 was derived from Compound 1 and Y-27632, we predicted it would be a competitive inhibitor of ATP binding to PKCε. Therefore, we assayed PKCε activity with increasing concentrations of ATP and in the presence of increasing concentrations of CP612. The EC50 for ATP was 4.2 µM (2.23 to 7.61 µM, 95% C.I.) and it increased 40-fold to 168 µM (121 to 250 µM, 95% C.I.) in the presence of 300 nM CP612 (Figure 1B). This rightward shift indicates that CP612 inhibits PKCε by competing with ATP.

We next performed a pharmacokinetics study in rats, which showed that CP612 (40 mg/kg) has a plasma half-life of 6.8 h ± 0.7 h after i.p. administration and of 4.0 h ± 0.6 h after i.v. administration (Figure 2A). The C_max_ was 29.3 ± 1.0 µM between 30 and 60 minutes after i.p. administration and 104.5 ± 3.1 µM at the earliest timepoint (5 min) after i.v. administration. Extrapolating back to time = 0, the initial i.v. concentration would be expected to be 110 µM. Despite significantly decreased plasma concentrations in the i.v. group between 2 and 24 hours, brain levels decreased less than 2-fold (from 2.6 µM to 1.5 µM, Figure 2A) resulting in an increase in the brain/plasma concentration ratio from 16 ± 3% at 2h to 917 ± 319% at 24h after injection. Plasma and brain levels were lower after i.p. administration (Figure 2B) but again brain levels remained stable between 2 and 24h (∼0.5 µM, Figure 2B), resulting in an increase in the brain/plasma concentration ratio from 3 ± 0.9% at 2h to 118 ± 42% at 24h after injection. These results indicate that CP612 enters the brain and is cleared from brain at a much slower rate than from plasma.

**Figure 2.**
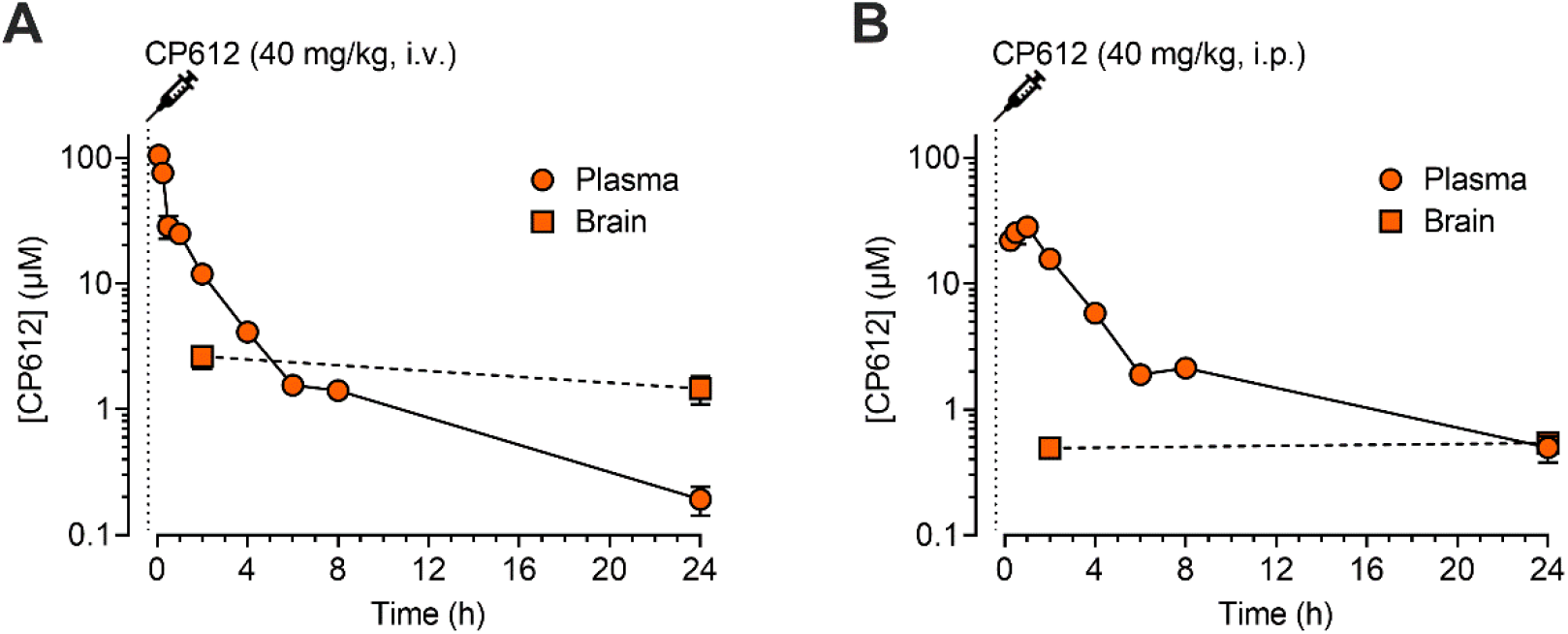
Pharmacokinetics of CP612. Concentrations of CP612 (40 mg/kg) in male mice after **(A)** i.v. or **(B)** i.p. administration declined over time in plasma but remained more stable in brain up to 24 h after administration. Data are mean and SEM (n = 2-3 per group).

### CP612 reduces PKCε-dependent hyperalgesia

To directly test if CP612 inhibits PKCε-dependent hyperalgesia, we administered the highly selective PKCε activator, ψεRACK, into the dorsum of the hind paw (1 µg/5 µL, intradermally) of rats, at the site of nociceptive threshold testing, to induce mechanical hyperalgesia. Thirty minutes later, rats were administered vehicle control (5 µL, intradermally) or CP612 (1 µg/5 µL, intradermally) (Figure 3). We measured nociceptive thresholds at baseline, 30 minutes after the administration of ψεRACK (time 0), and 15, 30, and 60 min, after the administration of CP612 or vehicle. Pain thresholds at baseline and at time 0 did not differ across groups (p values > 0.999). ψεRACK induced hyperalgesia in both groups (p values < 0.001) and this was reduced by CP612 at all timepoints tested [Figure 3, F _PKC inhibition_ (1,10) = 78.74, p < 0.001; F _Time x PKC inhibition_ (6, 60) = 7.29, p < 0.001]. Given the near complete reversal, these experiments indicate that CP612 inhibits PKCε-dependent mechanical hyperalgesia.

**Figure 3.**
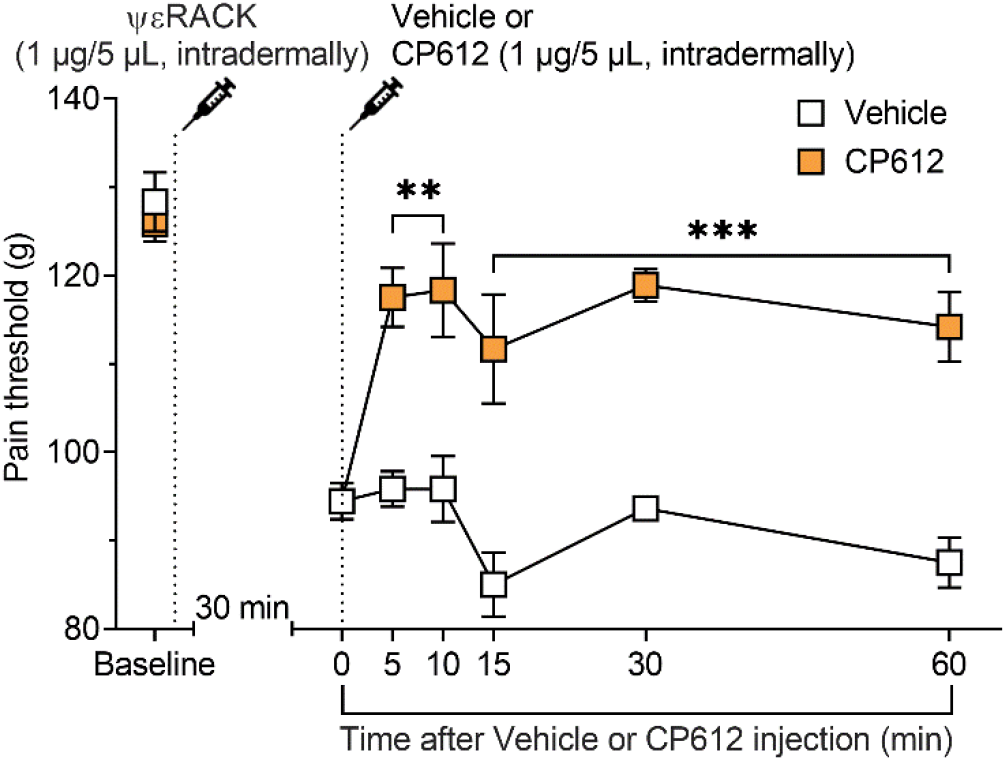
PKCε-induced hyperalgesia. The PKCε activator ψεRACK (1 µg/5 µL, intradermally) induced hyperalgesia in rats. This was attenuated by CP612 (1 µg/5 µL, intradermally), up to 60 min after its administration. Data are mean and SEM (n = 6 paws per group). ***p < 0.001 compared with the same timepoints in the Vehicle group.

### CP612 reduces paclitaxel-induced hyperalgesia

Repeated administration of the cancer chemotherapeutic drug paclitaxel produces long-lasting mechanical hyperalgesia (19). We tested if CP612 reverses this hyperalgesia. Male rats received repeated injections of paclitaxel (1 mg/kg, i.p. on days 0, 2, 4, and 6). Twenty-four hours after the last dose of paclitaxel, rats were administered vehicle control (2 mL/kg, i.p.) or CP612 (20 mg/kg, i.p.). We measured nociceptive thresholds at baseline, 24 h after the last injection of paclitaxel (time 0), and 15, 30, 60, 120 min and 24h after administration of CP612 or vehicle (Figure 4). Pain thresholds at baseline and at time 0 did not differ significantly across groups (p values > 0.946). Paclitaxel induced hyperalgesia in both groups (p values < 0.001), that was attenuated 15, 30, 60, and 120 minutes after the administration of CP612 [Figure 4, F _PKC inhibition_ (1,10) = 33.74, p < 0.001; F _Time x PKC inhibition_ (6,60) = 10.19, p < 0.0001]. This decrease in hyperalgesia dissipated one day after the administration of CP612. These results indicate that CP612 blocks paclitaxel-induced hyperalgesia and this effect dissipates after one day.

**Figure 4.**
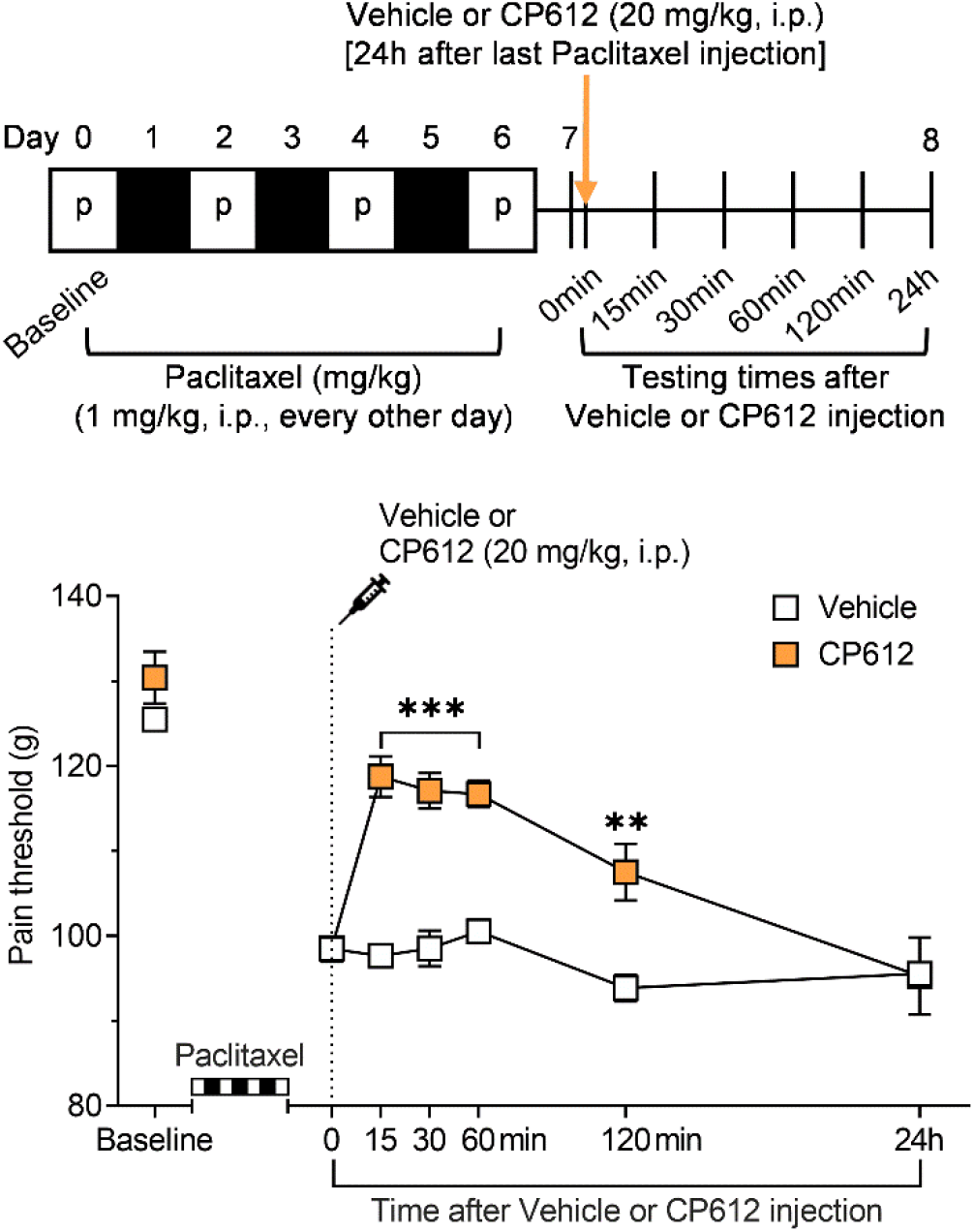
Paclitaxel-induced hyperalgesia. Repeated administration of paclitaxel (1 mg/kg, i.p. every other day for a total of 4 injections) induced hyperalgesia in male rats. This was attenuated by CP612, up to 120 min after its administration. Data are mean and SEM (n = 6 paws per group). **p < 0.01, ***p < 0.001 compared with the same timepoints in the Vehicle group.

### CP612 has a low “addictive potential”

We tested the addictive potential of CP612, by examining the extent to which rats are willing to work to obtain CP612. Rats were initially trained to self-administer either saline, morphine (500 µg/kg/infusion) or one of three different doses of CP612 (75 or 150 or 300 µg/kg/infusion) for four days at FR1 (fixed ratio 1, where one nose-pokes delivers one infusion). During this phase, rats learned self-administration behavior, as attested by greater responding in the active hole than the inactive hole [F _hole type_ (1,37) = 129.11, p < 0.001, data not shown]. Intake differed across different treatment groups [F _Group_ (4,37) = 8.50, p < 0.01) such that the group self-administering morphine took fewer infusions than all other groups (average intake 9.1 infusions), which had similar levels of intake (average intake 28.0, 29.3, 30.8, and 29.9 for rats self-administering saline or CP612 at 75, 150, or 300 µg/kg/infusion, respectively).

During the second phase, the ratio to obtain an infusion increased geometrically every other day (FR1, 3, 6, 12, 24….). The increase in ratio produced an increase in responding in the active hole [Figure 5A, F _Ratio_ (6,222) = 7.75, p < 0.001], and this occurred differentially across groups [F _Ratio x Group_ (24,222) = 15.71, p < 0.001]. Thus, rats self-administering saline or any dose of CP612 did not significantly increase responding in the active hole when the ratio to obtain an infusion increased, whereas rats self-administering morphine showed increased responding with the increase in ratio (Figure 5A). There were no group differences in responding in the inactive hole [F _Group_ (4,37) = 1.40, p > 0.25].

**Figure 5.**
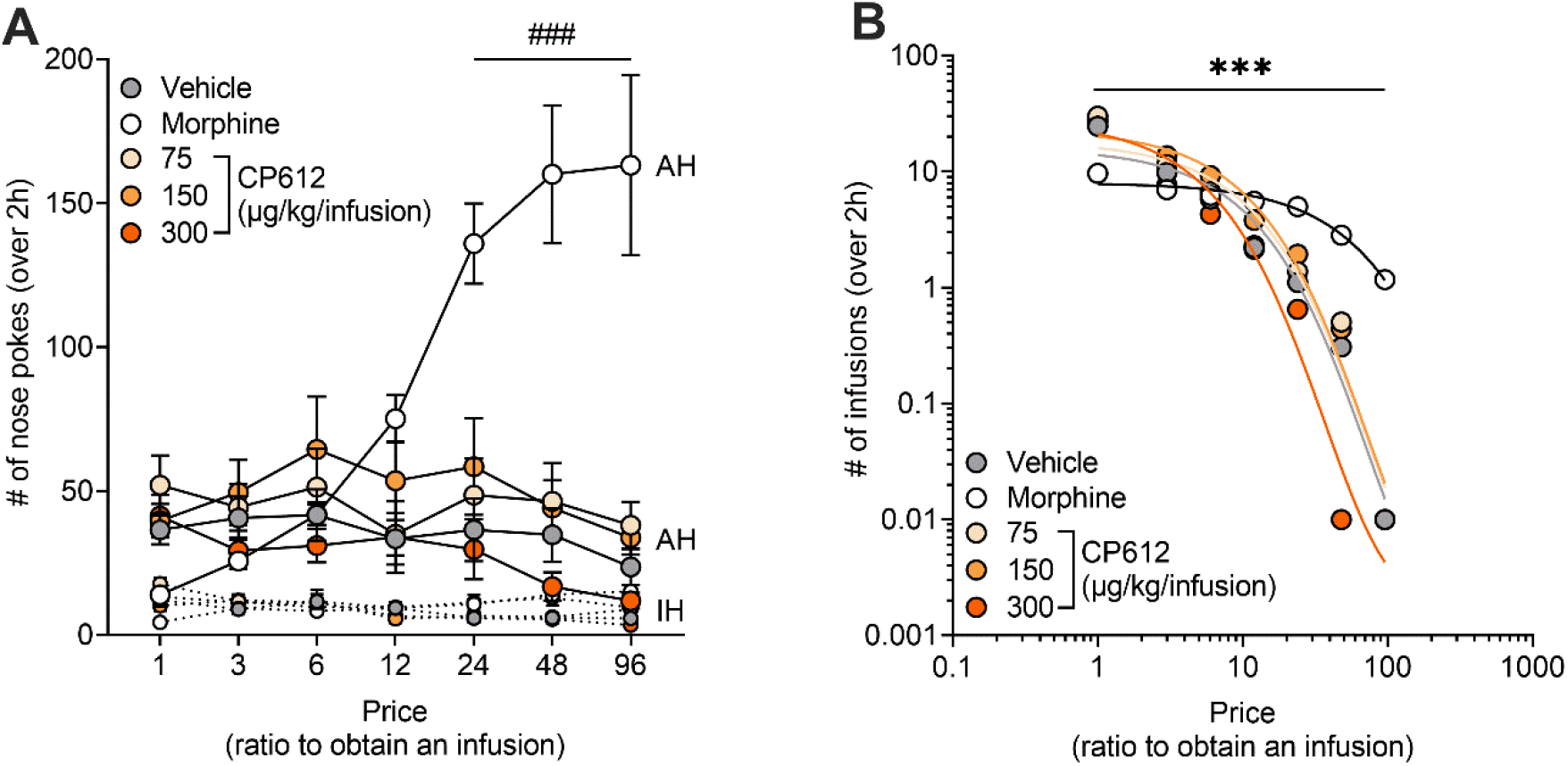
Self-administration of CP612. **(A)** Increasing the ratio to obtain an infusion across self-administration sessions produced a concomitant increase in responding in the active hole in rats self-administering morphine (500 µg/kg/infusion) but not in rats self-administering saline or different doses of CP612 (75, 150, and 300 µg/kg/infusion). **(B)** Rats self-administering morphine had a lower elasticity value (α = 0.4 × 10^3^) compared with all other groups [α = 1.1, 0.9, 0.9, 1.2 × 10^3^ for vehicle and CP612 at 75, 150, and 300 µg/kg/infusion]. They also had a higher price value to switch from inelastic to elastic behavior (Pmax = 50.1) compared with all other groups (Pmax = 4.1, 6.5, 6.1, 2.5 for vehicle and 75, 150, and 300 µg/kg/infusion). Data are mean and SEM (n = 10 for vehicle, n = 9 for morphine, n = 7-8 for each CP612 group). ###p < 0.001 compared with their infusions at FR1 and the same ratios in all other groups. Pmax: ***p < 0.001 morphine compared with all other groups. AH: Active hole; IH: Inactive hole.

The increase in ratio also changed the number of self-infusions [F _Ratio_ (6,222) = 171.68, p < 0.001], and this occurred differentially across groups [F _Ratio x Group_ (24,222) = 6.49, p < 0.001]. Thus, rats self-administering saline or any dose of CP612 showed a decrease in the number of self-infusions when the ratio to obtain an infusion increased. In contrast, rats self-administering morphine did not statistically decrease the number of self-infusions until FR98.

Data (number of self-infusions) were also fitted in an economic-demand curve, which can estimate the motivation to obtain drugs and their “addictive potential” (20). We measured elasticity (α), which is the relative change in consumption as a function of price (i.e. ratio of nose pokes required to obtain an infusion). Behavior is considered “inelastic” when consumption is insensitive to price and “elastic” when consumption is sensitive to price. We also measured Pmax, which is the price at which intake switches from inelastic to elastic. Rats self-administering morphine had a lower elasticity (α) compared with all other groups [F _Group_ (4,37) = 5.12, p < 0.01], and greater Pmax [F _Group_ (4,37) = 18.67, p < 0.001] (Figure 5B). Taken together, these findings indicate that CP612 is highly unlikely to be addictive.

### CP612 does not alter morphine self-administration

Because many patients with chronic pain take opioids for pain management, we wanted to know if CP612 would change opioid intake. Therefore, we investigated if adding CP612 to the morphine solution during self-administration (i.e., a cocktail of morphine and CP612) modifies self-administration of morphine. We tested rats from the prior experiment for self-administration of morphine (500 µg/kg/infusion). Rats were first tested under a FR6 to re-establish responding. The, we added vehicle or three different doses of CP612 (75,150, or 300 µg/kg/infusion) to the morphine solution. We examined its effects during a 4h session using a within-session progressive ratio schedule of reinforcement, in which the ratio to obtain an infusion is increased semi-logarithmically within a self-administration session (1, 2, 4, 6, 9, 12, 15, 20, 25, etc.), thereby allowing testing over a single day (21). CP612 did not modify responding at any dose tested, measured as number of infusions and breaking point (the highest ratio rats complete to earn an infusion of drug) [Figure 6, F _PKC inhibition_ (3,29) = 1.67, p > 0.195 and = 1.21, p > 0.324 for infusions and breaking point, respectively].

**Figure 6.**
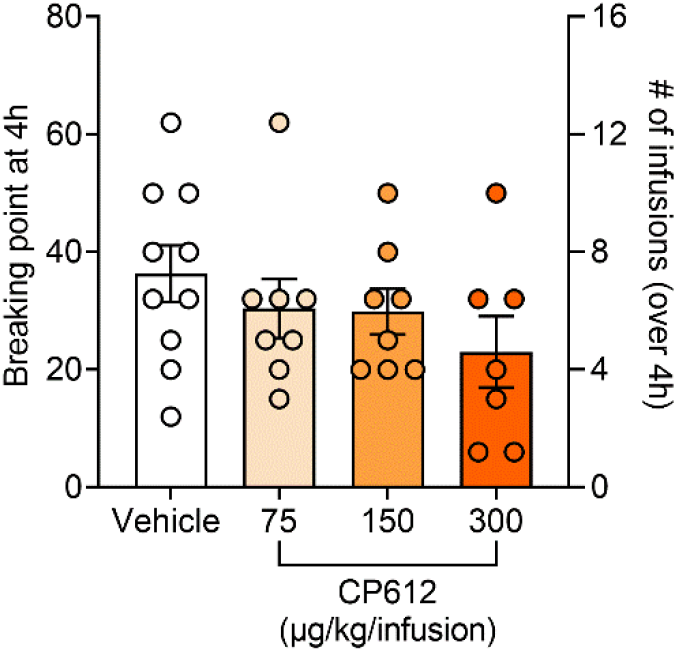
CP612 administered with morphine does not alter morphine self-administration. Adding CP612 (75, 150, and 300 µg/kg/infusion) to the morphine solution (500 µg/kg/infusion) during self-administration did not modify responding in a progressive ratio test, where the ratio to obtain an infusion was increased progressively within a self-administration session. This was measured as breaking point (highest ratio reached to earn an infusion of drug) and number of self-infusions. Bars are mean and SEM; dots are scores from individual rats (n = 7-10 per group).

We next investigated if CP612 administered prior to a self-administration session modifies self-administration of morphine. A separate group of rats was first trained to self-administer morphine (500 µg/kg/infusion) at FR1 and then FR3. Rats learned self-administration behavior, as attested by greater responding in the active hole vs. the inactive hole [F _Hole type_ (1,17) = 173.96, p < 0.001]; this occurred to a similar extent in rats that would later receive vehicle or CP612 [Figure 7A, F _Hole type x PKC inhibition_ (1,17) = 0.08, p > 0.78]. The average intake of morphine was 9.26 infusions, and this intake was similar in rats that would later receive vehicle or CP612 [F _PKC inhibition_ (1,17) = 0.02, p > 0.89; F _Days x PKC inhibition_ (8,126) = 0.65, p > 0.73].

**Figure 7.**
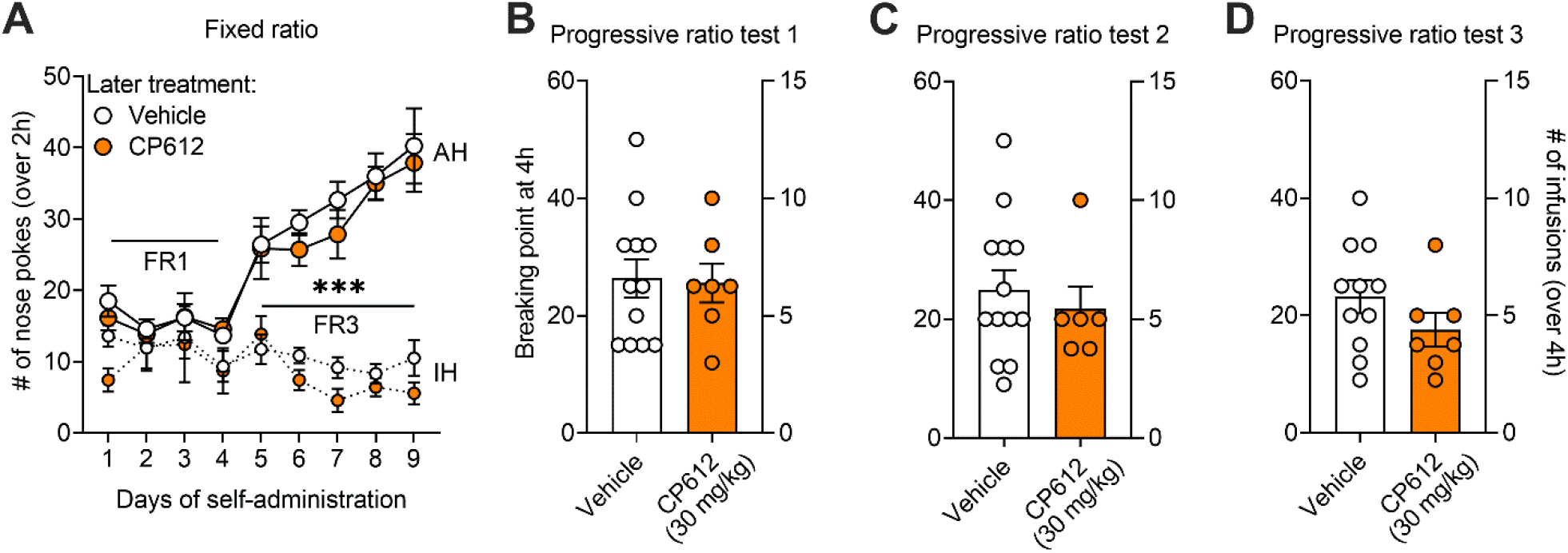
CP612 does not alter subsequent morphine self-administration. **(A)** Rats acquired morphine self-administration behavior at FR1, and this responding was increased when the ratio to obtain morphine was increased to FR3. This occurred to the same extent in rats that would later receive vehicle or CP612 (AH: Active hole; IH: Inactive hole). During the progressive ratio tests, the ratio to obtain an infusion was increased progressively within a self-administration session. Administration of CP612 (30 mg/kg, i.p.) 6h prior did not modify morphine intake or responding. This was measured as breaking point (highest ratio reached to earn an infusion of drug) and number of self-infusions **(B)** the day after the last self-administration session, **(C)** after one day of withdrawal or **(D)** after the administration of naloxone (0.03 mg/kg, s.c.). Data are mean and SEM; dots are scores from individual rats (n = 6-13 per group). ***p < 0.001 compared with the inactive hole.

CP612 (30 mg/kg, i.p.), administered 6h prior, did not modify morphine intake or responding, measured as number of morphine infusions and breaking point (the highest ratio rats complete to earn an infusion of morphine); this was true for all successive progressive ratio tests: test 1, done the day after initial self-administration [Figure 7B, F _PKC inhibition_ (1,17) = 0.01, p > 0.94 and = 0.02, p > 0.88 for infusions and breaking point, respectively], test 2, done after one day of withdrawal from self-administration [Figure 7C, F _PKC inhibition_ (1,17) = 0.16, p > 0.69 and = 0.34, p > 0.56 for infusions and breaking point, respectively], and test 3, done after the administration of naloxone (0.03 mg/kg, s.c.) [Figure 7D, F _PKC inhibition_ (1,17) = 1.66, p > 0.219 and = 1.80, p > 0.19 for infusions and breaking point, respectively]. Similar results were obtained when CP612 was administered 18h prior to the self-administration session (Figure S1).

### CP612 prevents and reverses hyperalgesia induced by morphine withdrawal

Many patients with chronic pain receive long-term opioid therapy, which can become ineffective over time and difficult to taper since reducing the dose of opioid medications can worsen pain (22). This effect can be modeled in rodents, where withdrawal from repeated administration of morphine produces hyperalgesia that is long lasting (23). To examine if CP612 alters this hyperalgesia, we first identified the timepoints at which hyperalgesia develops in male C57BL/6J mice. Mice received repeated injections of saline (10 mL/kg, i.p.) or morphine (20-100 mg/kg, i.p.) twice daily for five days, to induce dependence. Withdrawal from repeated administration of morphine induced hyperalgesia in a time-dependent manner [Figure 8, F _Opioid treatment_ (1,10) = 7.85, p < 0.05; F _Time x Opioid treatment_ (3,30) = 10.67, p < 0.001]. Specifically, hyperalgesia was not present at 6h but was present at 24h and 1 week after the last injection of morphine. Repeated administration of saline did not alter pain thresholds from baseline at any time tested.

**Figure 8.**
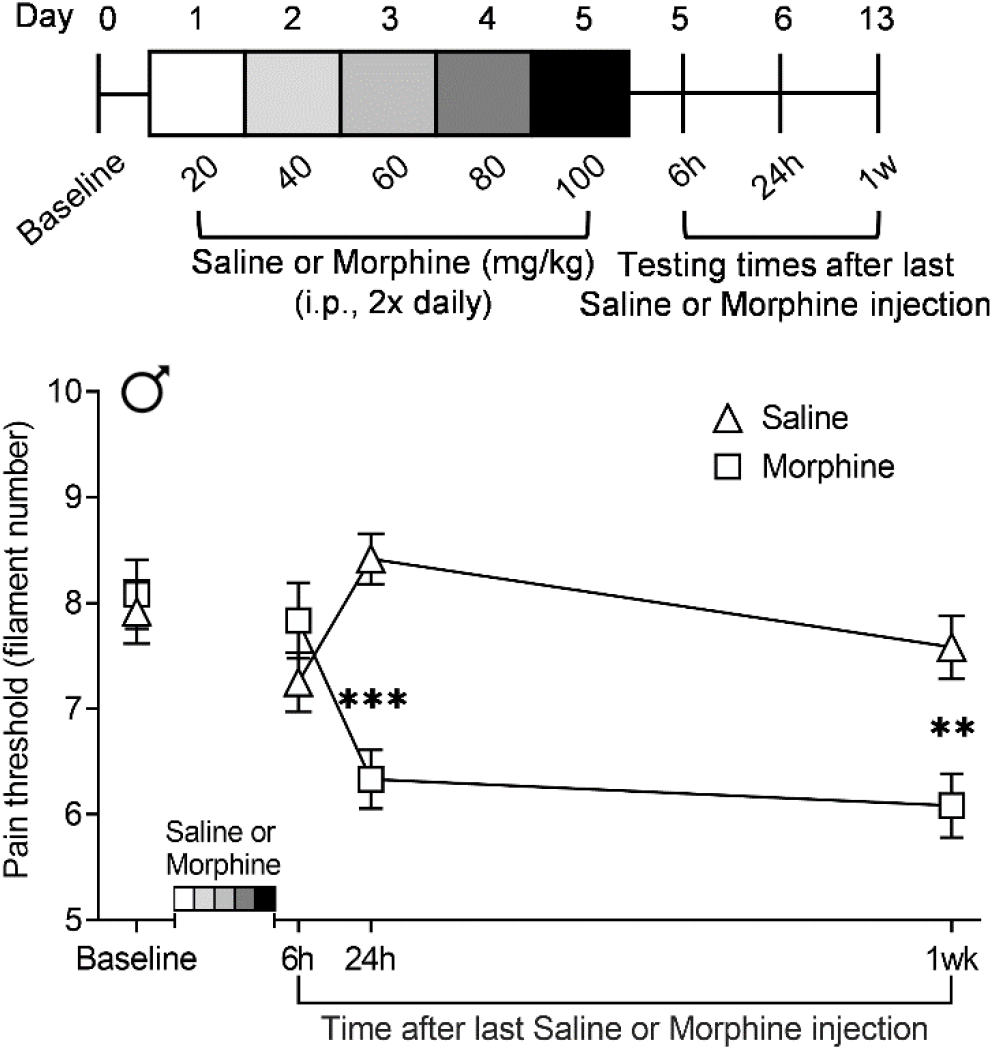
Time course of hyperalgesia induced by morphine withdrawal. After repeated administration of morphine (20-100 mg/kg, i.p.) twice daily for 5 days in male mice, hyperalgesia was not present at 6h but was present at 24h after the last injection of morphine and persisted for 1 week. No hyperalgesia developed after repeated administration of saline. Data are mean and SEM (n = 6 per group). **p < 0.01, ***p < 0.001 compared with the same timepoints in the group receiving repeated injections of saline.

We next investigated if CP612 *prevents* hyperalgesia induced by withdrawal from repeated administration of morphine, in male and female C57BL/6J mice. Mice received repeated injections of saline (10 mL/kg, i.p.) or morphine (20-100 mg/kg, i.p.) twice daily for five days. Six hours after the last injection (i.e., prior to the development of hyperalgesia), mice were administered vehicle control (10 mL/kg, i.p.) or CP612 (20 mg/kg, i.p.). Baseline pain thresholds did not differ significantly before treatment with saline, morphine, vehicle, or CP612 (p values > 0.99). Withdrawal from repeated administration of morphine produced hyperalgesia that was prevented by administration of CP612 whereas withdrawal from repeated administration of saline did not produce hyperalgesia and was not modified by administration of CP612 [Figure 9A-B, F _Opioid treatment_ (1,64) = 50.85, p < 0.001; F _PKC inhibition_ (1,64) = 52.08, p < 0.001; F _Opioid treatment x PKC inhibition_ (1,64) = 51.50, p < 0.001]. Results did not differ across sexes [F _Sex x Opioid treatment x PKC inhibition_ (1,64) = 2.34, p > 0.131; F _Time x Sex x Opioid treatment x PKC inhibition_ (4,256) = 1.23, p > 0.297). These findings indicate that a single dose of the PKCε inhibitor can prevent hyperalgesia induced by opioid withdrawal.

**Figure 9.**
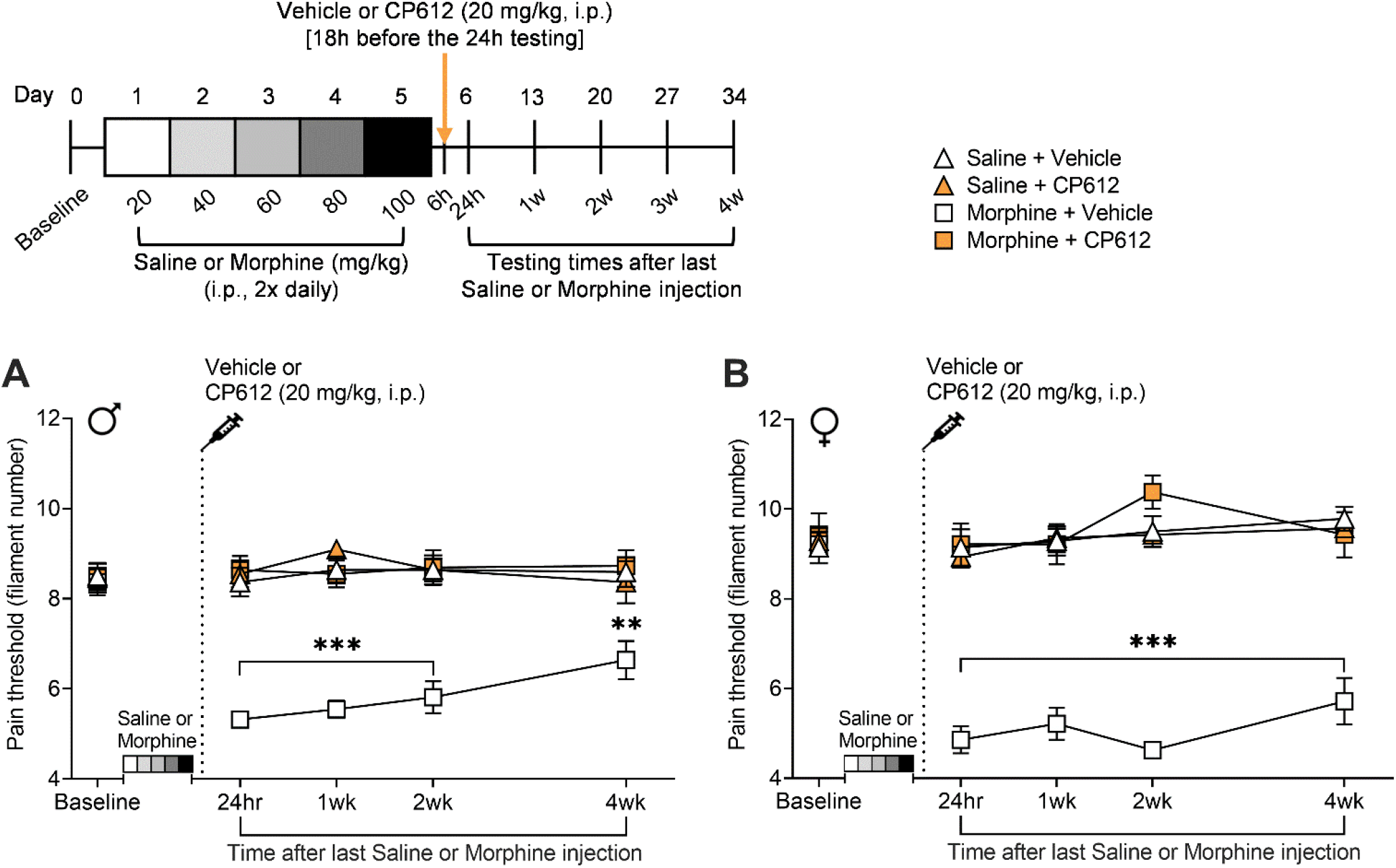
Prevention of hyperalgesia due to morphine withdrawal. Withdrawal from repeated administration of morphine (20-100 mg/kg, i.p.) induced hyperalgesia in **(A)** male and **(B)** female mice. This was prevented by CP612, up to 4 weeks after its administration. No hyperalgesia developed after repeated administration of saline. Data are mean and SEM (n = 11 for each male group, n = 7 for each female group). ***p < 0.001 and ** p < 0.01 compared with the same timepoints in all other groups.

We next tested if CP612 *reverses* hyperalgesia induced by withdrawal from repeated administration of morphine. Since the pain thresholds of mice receiving repeated administration of saline were not affected by treatment with vehicle or CP612 injection (Figure 9), mice in this experiment were only treated with morphine. Male and female C57BL/6J mice received repeated injections of morphine (20-100 mg/kg, i.p.) twice daily for five days. Two weeks after the last injection of morphine (i.e., after the development of hyperalgesia), mice were administered vehicle control (10 mL/kg, i.p.) or CP612 (20 mg/kg, i.p.). Baseline pain thresholds did not differ significantly before repeated administration of morphine (p values > 0.373). Hyperalgesia was apparent at 24h, and 1 week after the last injection of morphine. CP612 reversed the hyperalgesia [Figure 10A-B, F _PKC inhibition_ (1,18) = 53.53, p < 0.001], and this occurred similarly across sexes [F _sex x PKC inhibition_ (1,18) = 2.46, p > 0.133] and time [F _time x sex x PKC inhibition_ (5,90) = 0.23, p > 0.947]. Both male and female mice that received vehicle continued to show hyperalgesia up to 4 weeks after the last injection of morphine. Instead, mice that received CP612 showed a persistent reversal of hyperalgesia. These results indicate that a single dose of the PKCε inhibitor can reverse hyperalgesia induced by opioid withdrawal.

**Figure 10.**
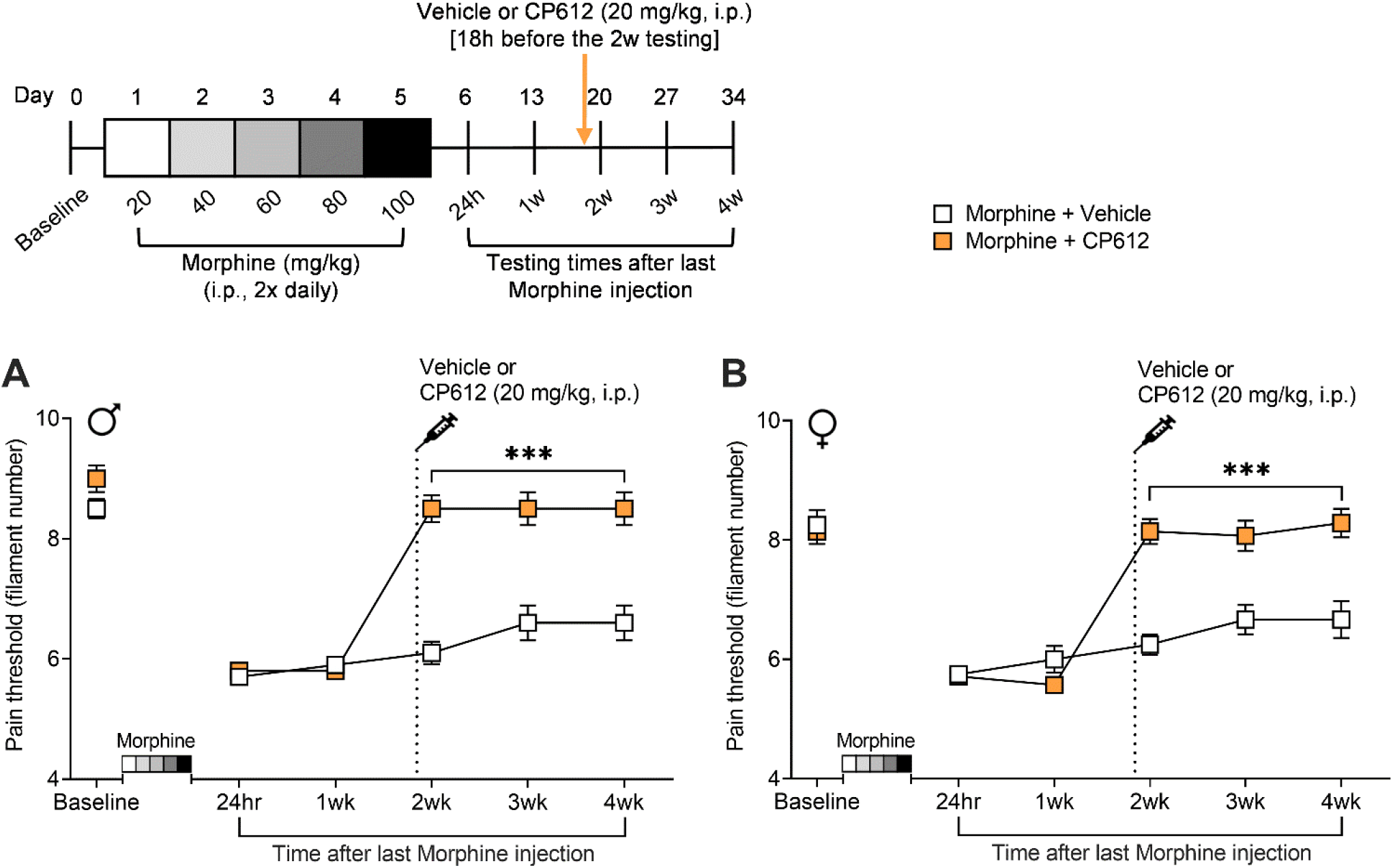
Reversal of hyperalgesia due to morphine withdrawal. Withdrawal from repeated administration of morphine (20-100 mg/kg, i.p.) induced hyperalgesia in **(A)** male and **(B)** female mice. This was reversed by CP612, administered 2 weeks after hyperalgesia was established and lasted for an additional two week. Data are mean and SEM (n = 5-7 per group). ***p < 0.001 compared with the same timepoints in the Vehicle group.

### CP612 reduces somatic signs of morphine withdrawal but not conditioned place aversion

In addition to hyperalgesia, withdrawal from repeated administration of morphine produces somatic signs of withdrawal and conditioned place aversion. We tested the effects of CP612 on these responses. A pilot experiment using C57BL/6 mice showed poor conditioned place aversion, so we used DBA/2J mice for these experiments, as they show more robust conditioned place aversion (24). Male DBA/2J mice received repeated injections of saline (10 mL/kg, i.p.) or morphine (20-100 mg/kg, i.p.) twice daily for 5 days. On day 6 (the conditioning day), mice were pretreated with vehicle control (10 mL/kg, i.p.) or CP612 (different doses, i.p.), followed by a single injection of saline (10 mL/kg, i.p.) or morphine (100 mg/kg, i.p.) 4h later. Two hours after this last injection of saline or morphine, mice were administered naloxone (5 mg/kg, i.p.) to precipitate somatic signs of withdrawal and conditioned place aversion. Withdrawal from repeated administration of morphine evoked somatic signs of withdrawal (Figure 11A and Figure S2); only the highest dose of CP612 (40 mg/kg) reduced them [Figure 11A, H _PKC inhibition_ (3, n = 62) = 33.34, p < 0.001]. Withdrawal from repeated administration of morphine also evoked conditioned place aversion, [Figure 11B, F _PKC inhibition_ (5,78) = 7.16, p < 0.001; mice receiving repeated administration of saline with vehicle vs. mice receiving repeated administration of morphine with vehicle, p < 0.001] and this was not modified by CP612 at any dose tested (p values > 0.079).

**Figure 11.**
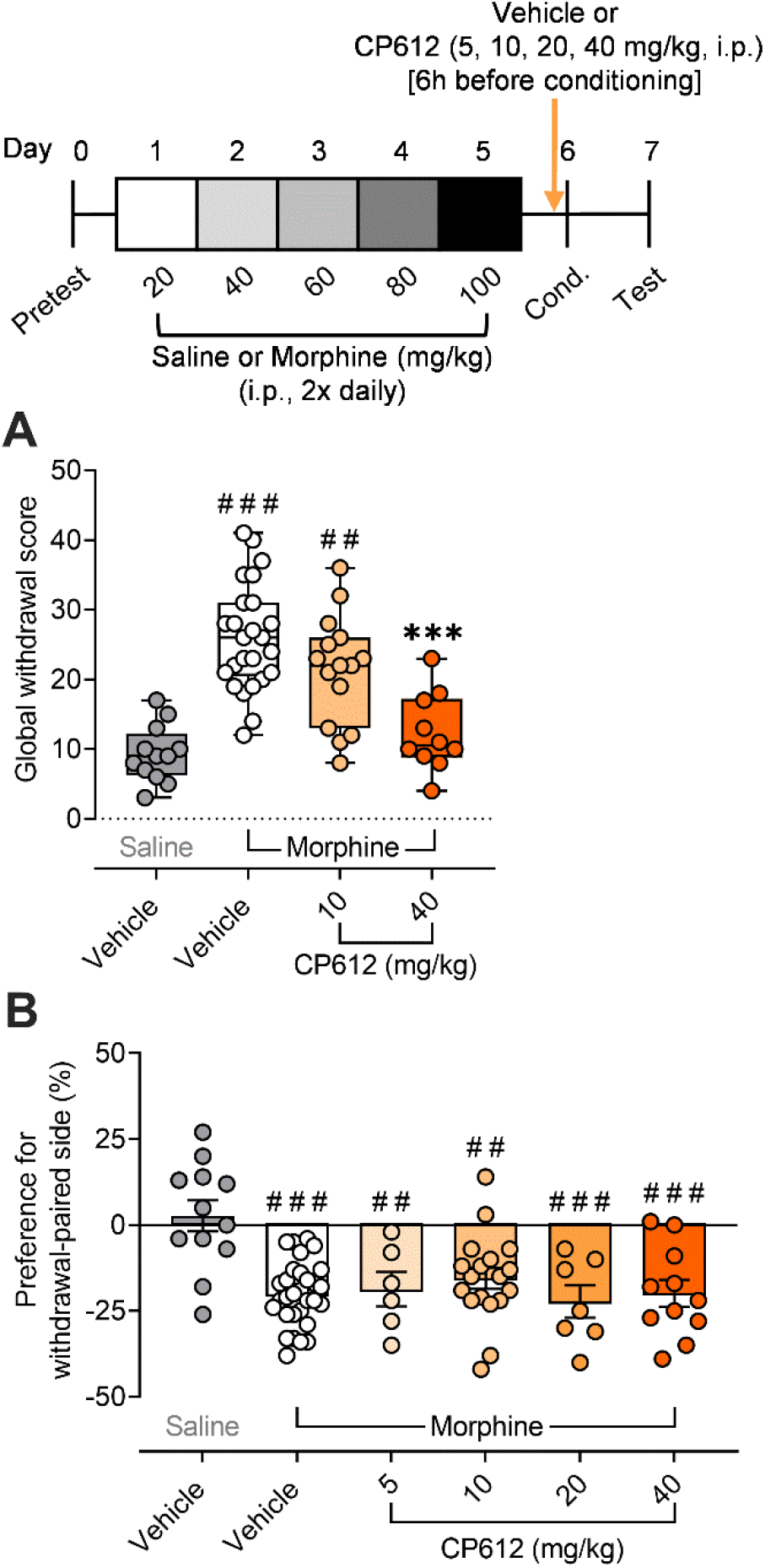
Morphine withdrawal and CPA precipitated by naloxone. **(A)** On conditioning day, naloxone (5 mg/kg, i.p.) evoked signs of withdrawal in male mice that received repeated administration of morphine (20-100 mg/kg, i.p.) and were treated with vehicle (n = 25) or a low dose of CP612 (10 mg/kg, i.p., n = 15), but this was decreased in mice treated with a high dose of CP612 (40 mg/kg, i.p., n = 10). **(B)** On test day, prior administration of naloxone produced CPA in mice that received repeated administration of morphine, and this occurred similarly in mice that were treated with vehicle (n = 30) or different doses of CP612 (5, 10, 20, 40 mg/kg, i.p., n = 6, 18, 7, 11, respectively). For (A), box and whiskers are median and 25% interquartile intervals; dots are scores from individual mice. For (B), bars are mean and SEM; dots are scores from individual mice. ##p < 0.01 and ###p < 0.001 compared with Saline + Vehicle: ***p < 0.001 compared with Morphine + Vehicle.

## Discussion

We describe a new small molecule inhibitor of PKCε, CP612, that reduces hyperalgesia evoked by activation of PKCε in peripheral nociceptors and by administration of paclitaxel in rats. CP612 itself had no addictive potential, nor did it modify the addictive potential of morphine but it prevented and reversed hyperalgesia induced by morphine withdrawal. This effect did not generalize well to other features of opioid withdrawal such as conditioned place aversion or somatic signs of withdrawal, which was reduced only at a high dose but not at a low dose of CP612 that inhibited pain.

In-vitro analysis showed that CP612 inhibited PKCε competitively with respect to ATP with low nanomolar potency. CP612 interacted only with ROCK1 and 2, novel PKC isozymes, and 5 other kinases when screened at 200 nM against a panel of 468 native kinases. CP612 was 40-fold more potent at inhibiting PKCε than ROCK1, and within the novel PKC subfamily, was most potent against PKCε. Pharmacokinetics studies in rats showed that CP612 was brain penetrant and had long CNS half-life when administered by intraperitoneal or intravenous injection. Together, these properties suggest that CP612 is a potent and selective inhibitor of PKCε that has favorable CNS drug-like physical-chemical properties based on those measured for clinically useful CNS drugs by CNS multiparameter optimization analysis (18).

Paclitaxel-induced hyperalgesia is long lasting in rats, persisting for up to 2 weeks after the last injection (19). Although CP612 reduced hyperalgesia in this model, the effect dissipated within 24h. We also found that hyperalgesia produced by withdrawal from repeated administration of morphine persisted in mice for over 1 week, as reported by others (23, 25, 26). CP612 abolished this hyperalgesia when administered before it had developed and later when it was well established. Unlike with paclitaxel-induced hyperalgesia, this effect of CP612 was persistent, lasting several weeks. These differences in response may reflect differences in pathophysiology between paclitaxel, which causes a toxic neuropathy, and morphine, which evokes a neuroadaptive response. That the effect of a single injection of CP612 was long-lasting, suggests that a single dose of a PKCε inhibitor could be useful to prevent or reverse hyperalgesia provoked by opioid withdrawal.

Previous studies have identified sexual dimorphism for PKCε in different models of chronic pain. Inhibition of PKCε was only able to attenuate hyperalgesia in male but not female rats with diabetic hyperalgesia induced by vincristine- or streptozotocin (27, 28). We found no sex differences in the effect of CP612 on prevention or reversal of hyperalgesia induced by opioid withdrawal. However, further work is needed to determine if there are sex differences in the effects of CP612 in paclitaxel-induced hyperalgesia since studies to date have only investigated the role of PKCε in male rats (19) and mice (29).

In addition to producing hyperalgesia, opioid withdrawal evokes an aversive state thought to promote opioid use (30, 31). Conditioned place aversion (CPA) is a sensitive index of this aversive motivational state and can be tested in rodents (32). We found that CP612 (5-40 mg/kg) did not block CPA induced by opioid withdrawal in mice. Despite not affecting CPA, CP612 had a dose-dependent effect on somatic signs induced by opioid withdrawal. A low dose of CP612 (10 mg/kg) had no effect, but a higher dose (40 mg/kg) reduced somatic withdrawal signs. These differences likely reflect selective actions of PKCε in pain pathways versus reward and other circuits that mediate somatic signs of withdrawal.

When developing new pain medications, it is important to determine if they have rewarding properties themselves, thereby posing an addiction risk. We tested this possibility by comparing self-administration of CP612 to self-administration of morphine or saline. This was done using a between-session progressive ratio test, whereby the ratio to obtain the drug increases geometrically across successive days. This allows fitting data to an economic demand-curve, that can estimate the motivation to obtain drugs and their “addictive potential” (20). Rats were initially tested at a fixed ratio 1 (one nose-poke yields one infusion of drug). During this phase, self-administration of morphine was low, as is expected at this dose (500 µg/kg/infusion) (33, 34). Instead, self-administration of saline was higher, and likely occurred because nose-poking in the active hole was coupled with the presentation of a light cue, which is reinforcing. Self-administration of saline occurred to the same extent as any dose of CP612 tested. As the ratio to obtain infusions increased, however, responding increased for morphine, but not for saline or any dose of CP612. In fact, the maximum price rats were willing to pay for an infusion of drug was highest for morphine than for all other groups, which did not differ from each other. Furthermore, intake of morphine was “inelastic” (did not change as the price to obtain the drug increased) compared with intake of saline and any dose of CP612, which was “elastic” (decreased as the price to obtain the drug increased). These results indicate that CP612 is unlikely to be addictive.

Because non-opioid pain medications might be taken with opioids, we tested if adding CP612 to the morphine solution during self-administration could enhance self-administration of morphine. CP612 did not modify morphine intake in a progressive ratio test at any dose tested, suggesting that CP612 is unlikely to increase motivation to self-administer morphine. We also tested if administering CP612 to rats that already learned to self-administer morphine could modify self-administration of morphine. Again, CP612 had no effect on morphine self-administration. These results suggest that CP612 is unlikely to modify morphine self-administration. A caveat of these experiments is that rats were trained to self-administer morphine at 500 µg/kg/infusion. This dose was chosen because it produces i.v. intake of approximately 5-7.5 mg/kg, which is within the range of i.v. doses that produce analgesia in rats (35). However, this dose does not produce dependence. It is therefore possible that, had rats self-administered higher doses of morphine leading to dependence, we might have observed different effects.

In conclusion, we have developed a non-addictive, small molecule inhibitor of PKCε, CP612 that was effective in treating pain in a model of chemotherapy-induced neuropathy. CP612, had the added and important benefit of preventing and reversing hyperalgesia evoked by opioid withdrawal. Long-term opioid therapy can become ineffective in controlling pain and improving function, and because opioid tapering can worsen pain, reducing the dose of opioid medications can be extremely difficult (22). Because CP612 was effective in preventing and reversing hyperalgesia due to opioid withdrawal, PKCε inhibitors may provide a new therapeutic option for opioid tapering in persons for whom risks associated with long-term opioid treatment outweigh the benefits.

## Materials and Methods

### Animals

Rats (Sprague-Dawley) were acquired from Envigo (Indianapolis, IN) and Charles River Laboratories (Hollister, CA) and housed 2-3 per cage. Male rats weighed 250-275 g and females weighed 175-199 g upon arrival.

C57BL/6J and DBA/2J mice were acquired from The Jackson Laboratory (Bar Harbor, Maine) and housed 2-3 per cage. Male and female mice were 8-9 weeks old and weighed 24-27 g (males) or 19-22 g (females) upon arrival.

Animals were housed on a 12:12 light:dark cycle with *ad libitum* access to water and laboratory chow (LabDiet, St. Louis, MO). Experiments were performed 8-10 hours after lights on, except for conditioned place aversion and self-administration experiments, which were performed 3-6 hours after lights off, to facilitate conditioning (36) and self-administration (37).

All experiments were performed on separate groups of animals unless specified.

### Drugs/reagents

Morphine sulfate was obtained from Spectrum Chemical (New Brunswick, NJ) and diluted in saline (0.9% NaCl). Doses are expressed as morphine base. Naloxone hydrochloride dihydrate was obtained from Spectrum Chemical (New Brunswick, NJ) and diluted in saline (0.9% NaCl). Mepivacaine was obtained from TCI America (Portland, OR) and diluted in saline (0.9% NaCl). Sodium brevital was obtained from Henry Schein (Dublin, OH) and was diluted in saline (0.9% NaCl). Lidocaine was obtained from Aspen Veterinary Resources (Liberty, MO) and diluted in saline (0.9% NaCl). The ψεRACK peptide (HDAPIGYD) was obtained from AnaSpec Inc. (Fremont, CA). CIDD-0150612 (hereafter called CP612) was synthesized in enantiomerically enriched form (> 96% enriched) by Dr. Stanton McHardy (University of Texas San Antonio, TX) and diluted in a vehicle solution made of 5% Tween 80 (Sigma Aldrich, Burlington, MA) and saline (0.9% NaCl) for i.p. injections and in saline (0.9% NaCl) for intravenous infusions.

### Kinase assays

CP612 was assayed against His-AVI-tagged human PKCε (1 nM) produced in Sf9 cells, and ROCK1 (Carna Biosciences, Natick, MA; 5 nM) using the Lance-FRET kinase assay system (PerkinElmer, Waltham, MA). Other human recombinant PKCs (SignalChem Biotech, Richmond, BC, Canada) were assayed in triplicate at the following concentrations: PKCβII 1.6 nM, PKCγ 500 pM, PKCδ 5 nM, PKCθ 500 pM, and PKCζ 500 pM. PKCs were incubated in a 15 µL reaction volume with kinase buffer (20 mM Tris-HCL, pH 7.4, 10 mM MgCl_2_, 0.25 mM EGTA, 0.1 mg/mL BSA and 0.15% TritonX-100), 50nM *ULight*™-labeled CREBtide CKRREILSRRPSYRK (PerkinElmer, Waltham, MA) and test compound. Reactions were initiated by adding 2.5 µM ATP at 27°C and were terminated after 60 min with 10 µL of 2.5X Stop Solution/Detection Mix containing 20mM EDTA and 4nM LANCE *Ultra* Europium-anti-phospho-PKC (Ala25Ser) peptide antibody (Perkin Elmer, Waltham, MA). After incubation at 27°C for another 60 min, phosphorylation was detected using a FlexStation 3 Microplate Reader (Molecular Devices, Sunnyvale, CA) in LANCE TR-FRET mode (excitation = 340 nm, emission = 665 nm) and expressed as relative fluorescence units (RFU). The percentage of inhibition was calculated as: (signal without test compound − signal with test compound) / (signal without test compound − signal without ATP) × 100. The Z’ factor for this assay ranged between 0.72 and 0.86.

### Pharmacokinetic studies in rats

CP612 was dissolved at 8 mg/mL in 10% Tween-80 / 90% water and administered at 40 mg/kg. Six rats were injected i.v. and six i.p. Three rats were used to collect a full plasma time course from 5 min to 24 hours post-dose. Then brains were collected from these rats at 24h and from three additional rats at 2h after administration of compound. Plasma samples were diluted 10-fold in blank plasma. Each brain was disrupted in potassium phosphate buffer (pH 7.4; 9 µL per 1mg tissue) on ice using brief sonication pulses. CP612 was detected with an API Triple Quad 5500 LC-MS/MS (SCIEX, Redwood City, CA) using an MRM(+) method: CP612 (463/282, m/z) with carbamazepine (238/195.1, m/z) as an internal standard.

## Mechanical nociception

### Rats

Mechanical nociceptive thresholds in rats were quantified by the Randall-Selitto paw withdrawal test (38) using an Ugo Basile Analgesiometer (Stoelting, Wood Dale, IL). This device applies a linearly increasing mechanical force to the dorsum of the rat hind paw. The mechanical nociceptive threshold is defined as the force in grams at which a rat withdraws its paw. The higher the force to elicit paw withdrawal, the higher the pain threshold (and the lower the sensitivity to pain). Rats were placed into cylindrical acrylic restrainers for 40 min before starting each session. The restrainers had lateral ports to allow access to the hind paw, as previously described (39). Prior to starting experiments, baseline mechanical nociceptive threshold was obtained as the mean of 3 readings on one hind paw in each rat.

### Mice

Mechanical nociceptive thresholds in mice were quantified using the simplified up-down (SUDO) method with von Frey filaments (40). The testing apparatus consisted of an elevated platform (34 × 72.5 × 35 in cm) with a metal mesh floor with square plastic compartments (10 × 14 × 10 in cm) as covers. von Frey filaments (Semmes-Weinstein monofilaments; Stoelting, Wood Dale, IL) were pressed against the center of the plantar surface of the hind paw until they buckled and were held for a maximum of 5 s. The force to bend the filaments ranged from 0.008 g (filament 1) to 6 g (filament 12). Mice were acclimated to the test room and apparatus for 1h prior to testing. We started testing mice with filament 5 (bending force 0.16 g) and stepped up or down according to the response of the mouse. The filament that elicited a definite paw withdrawal or notable flinch was recorded. All measures were averaged between both hind paws and are expressed as filament number, which allows parametric analysis. The higher the filament number the higher the pain threshold (and the lower the sensitivity to pain). Prior to starting experiments, baseline threshold levels were obtained 3-4 times at 2-5-day intervals on each of the hind paws of the mouse. Mice were then distributed in each experimental group according to their baseline level of response, to ensure similar baseline values across groups.

## Intravenous self-administration

### Implantation of intravenous catheters

Rats were anesthetized to implant catheters for intravenous self-administration. Anesthesia was induced by placing rats into an induction chamber filled with 5% isoflurane gas and was maintained with a vaporizer by delivering 2–3% isoflurane *via* a nose cone (E-Z Anesthesia, Palmer, PA). To ensure sufficient anesthesia, breathing rate, pinch response, and body temperature were monitored throughout surgery, and anesthesia was adjusted when necessary. Areas around incisions were shaved with electric clippers (Andis Company, Sturtevant, WI), and were cleaned with 70% alcohol, followed by 10% betadine and Lanacaine (20% benzocaine, 0.2% benzethonium chloride, 36% ethanol). Incision sites were then infused with the local anesthetics mepivacaine (2%) for the back incision and lidocaine (0.5%) for the front incision. Silastic catheters were implanted in the right external jugular vein and passed under the skin to exit in the mid-scapular region. The catheters were accessible through a backport that was mounted onto a pedestal secured under the skin with surgical staples (Braintree Scientific, Inc., Braintree, MA). At the conclusion of each surgery, wounds were covered with topical triple antibiotic ointment (bacitracin, neomycin sulfate, polymyxin B sulfate), (Medique Products, Fort Myers, FL). Systemic NSAID analgesics, either carprofen (5 mg/kg/mL, s.c.) or flunixin meglumine (2.5 mg/kg/0.5 mL, s.c.), were administered the day of surgery and 1-2 days following. Systemic antibiotic cefazolin (50 mg/kg/0.5 mL, i.v.) was administered the day of surgery and 2–6 days following, then reduced to 30 mg/kg/0.5 mL for the remainder of the experiment.

### Intravenous self-administration

Self-administration procedures took place in oversized Med Associates operant chambers (41 × 24 cm floor area, 21 cm high, Med Associates, Fairfax, VT) outfitted with photobeams to track locomotion, two nose-holes to track responding (nose-poking). Rats were placed daily in the operant chamber for 2-4h (depending on the study). The backport of the catheter was connected with a tether to an infusion pump (Med Associates, Fairfax, VT). The tether was made of Tygon tubing (Cole Parmer, Vernon Hills, IL) shielded by a metal spring (Instech Laboratories, Plymouth Meeting, PA), to prevent rats from chewing on it. We also added a “nylabone” in the chamber for chewing, as rats self-administering morphine often chewed on their tethers. Nose-poking into one hole (“active hole”) activated the infusion pump to deliver an intravenous infusion of drug at the rate of 11-12 µL/s and a volume of 150 µL/kg, concomitant with a 15s light cue within the hole and a 15s time-out period. Nose-poking into the other hole (“inactive hole”) had no consequences and was used to track non-goal-directed nose-poking. We recorded number of nose pokes and infusions using MED-PC IV software (Med Associates, Fairfax, VT). At the end of each experiment, we tested catheter patency by administering the fast-acting anesthetic sodium brevital (5 mg/kg/0.5 mL, i.v.). Rats not immediately anesthetized were eliminated from the study. For tests involving administration of CP612 or vehicle control after acquisition of self-administration of morphine, rats were distributed in each experimental group according to their baseline intake of morphine, ensure similar baseline values across groups.

### Progressive ratio testing (between-session)

Rats were tested for self-administration of saline, morphine (500 µg/kg/infusion), or three doses of CP612 (75 or 150 or 300 µg/kg/infusion) for 2h per day under a “fixed ratio 1” (FR1) schedule of reinforcement (1 nose poke in the “active hole” is required to obtain 1 infusion of drug) until behavior (number of infusions) was stable (varied less than 25% for 3 consecutive days); this lasted 4-7 days, depending on the rat. Following this initial acquisition phase at FR1, the ratio to obtain the drug was increased geometrically every other day: FR3 (3 nose pokes are required to obtain 1 infusion of morphine), 6, 12, 24, 48, 96. This allows analyzing drug consumption as a function of “price” using demand-curves for behavioral economics, which can estimate the motivation to obtain drugs and their “addictive potential” (20). For this, data from each rat were fitted to the equation log Q = log Q_0_ + k (e ^−α^(^Q0 C)^ – 1), where Q represents consumption (number of self-infusions), and Q_0_ the level of consumption at the lowest price. C represents price for the drug (i.e., the ratio), and k is set to a constant accounting for the estimated range of consumption in logarithmic units (k = 4 in these studies). α is the relative change in consumption across changes in price (ratio). Behavior is considered “inelastic” when consumption is insensitive to price (i.e., the curve is flat, and consumption is maintained despite increases in price) and it switches to “elastic” when consumption is sensitive to price (i.e., when the curve is steep and consumption declines with increases in price). A drug is deemed to have a higher addictive potential when it has low elasticity and when the switch from inelastic to elastic occurs at high prices (41, 42). Using this curve, we can also calculate Pmax, which is the price at which the switch from inelastic to elastic occurs and is calculated as 0.65/(Q_0_ × α × k^1.191^); greater Pmax indicate greater effort to obtain the reward (42). We measured responding in the active and inactive holes, number of self-infusions, and locomotor activity. We also compared groups for their average intake in the consumption-price curves by measuring α (elasticity) and Pmax (price at which the switch from inelastic to elastic occurs).

### Progressive ratio testing (within-session)

Rats were tested for self-administration in a within-session progressive ratio test in which the ratio to obtain one infusion increased semi-logarithmically (1, 2, 4, 6, 9, 12, 15, 20, 25, etc.) within a single 4-h self-administration session (21). This allows testing for the “breaking point” (the highest ratio reached to obtain one infusion).

#### Induction of morphine dependence

Morphine dependence was induced by repeated intraperitoneal (i.p.) injections (10 mL/kg) of morphine administered at increasing doses, over 5 days (day 1: 20 mg/kg, day 2: 40 mg/kg, day 3: 60 mg/kg, day 4: 80 mg/kg, day 5: 100 mg/kg). Injections of each dose were administered twice daily, 6h apart. This dosing was chosen because it produces spontaneous opioid withdrawal and withdrawal-induced mechanical hypersensitivity in mice (23).

#### Conditioned place aversion and somatic signs of morphine withdrawal

The place conditioning apparatus was an acrylic box enclosed in light- and sound-attenuating, ventilated chambers (MED Associates, Fairfax, VT). The box had two compartments separated by a removable sliding door. Compartment 1 had a floor comprised of metal bars (0.24 cm) placed in parallel series. Compartment 2 had a floor comprised of a metal grid mesh (0.63 cm). The dimensions of each compartment were 13.5 × 27 × 17 cm. Infrared light beams and photodetectors located along the walls of the box measured general activity and location of the mouse in each compartment which were recorded using Activity Monitor version 7.0.5.10 software (Med Associates, Fairfax, VT). On day 0 (pretest), drug-naïve mice were acclimated for 15 min to the place conditioning apparatus without the sliding door, so they had free access to both compartments of the apparatus. This established a baseline side-preference, based on the amount of time spent in each compartment. Mice that exhibited > 75% preference for one compartment were removed from the study. Then, mice were distributed in each experimental group according to their preference scores, to ensure similar values across groups. Then they received repeated injections of saline (10 mL/kg, i.p.) or morphine (20-100 mg/kg, i.p.), twice a day, for five days. On day 6 (conditioning day), mice were pretreated with vehicle (10 mL/kg, i.p.) or CP612 (5, 10, 20, or 40 mg/kg, i.p.), followed by a single injection of morphine (100 mg/kg, i.p.) 4h later. Two hours after this last injection of morphine, mice were administered naloxone (5 mg/kg, i.p.), to precipitate withdrawal, and were immediately confined to one side of the conditioning apparatus, for 20 min. During this conditioning period, mice were filmed (raspberry pi camera) to assess somatic signs of withdrawal (43). Given the limited number of cameras and the poor quality of some videos, we were only able to film a subset of mice. The following behaviors were given a score of 1 per occurrence and episode: body shakes (paw tremors and wet dog shakes), teeth chatter, rearing, hopping, wall climbing, and backpedaling. These scores were summed to obtain a global withdrawal score and were obtained from an independent observer who was blind to the experimental conditions. A subset of videos was scored by two observers, to ensure internal validity (T _Observer_ = 0.23, p > 0.827, Correlation _Observer_ r = 0.97, p < 0.005). On day 7 (test day), mice were placed in the conditioning apparatus with free access to both compartments, for 15 min. We recorded the time spent in each compartment.

#### Statistical analysis

Kinase assays used 11-point concentration-response curves and results were analyzed by nonlinear regression using Prism 9 (GraphPad Software) to determine IC50 values. Scores for pain threshold, conditioned place preference, and self-administration were analyzed by ANOVA using the following factors. Between-subject factors: PKC inhibition (vehicle, CP612 at different doses), chemotherapeutic treatment (vehicle, paclitaxel), opioid treatment (morphine, saline); Within-subject factors: Time (different times or days of testing), hole (active hole, inactive hole). We used a Tukey test for multiple comparisons for between-group comparison and Dunnett test for comparison with own baseline values in the pain studies. Somatic withdrawal scores were analyzed with a Kruskal Wallis test due to the non-parametric nature of those data, followed by non-parametric post-hoc comparisons according to Siegel and Castellan (44).

#### Study approval

All procedures were done following The National Institutes of Health Guide for the Care and Use of Laboratory Animals and were approved by the Institutional Animal Care and Use Committee of The University of Texas at Austin and of the University of California at San Francisco.

## Author contributions

ROM, JDL, SFM and MM conceived the study and designed the experiments. AG-F, IJMB, SD, HDK, NR, MM, and CF conducted the experiments. AG-F, IJMB, SD, HDK, and CF analyzed data in collaboration with ROM, JDL, SFM and MM. AG-F, ROM, JDL, SFM, and MM wrote the manuscript.

## Acknowledgements

This research was supported by the US Department of Defense Congressionally Directed Medical Research Programs (CDMRP) Peer Reviewed Medical Research Program (PRMRP) (W81XWH1810389 to ROM and JDL). Synthesis of CP612 used in these studies was supported under the Cancer Prevention Research Institute (RP210208 to SFM).

