## Supplementary figures for "A novel small molecule PKC epsilon inhibitor reduces hyperalgesia induced by paclitaxel or opioid withdrawal"

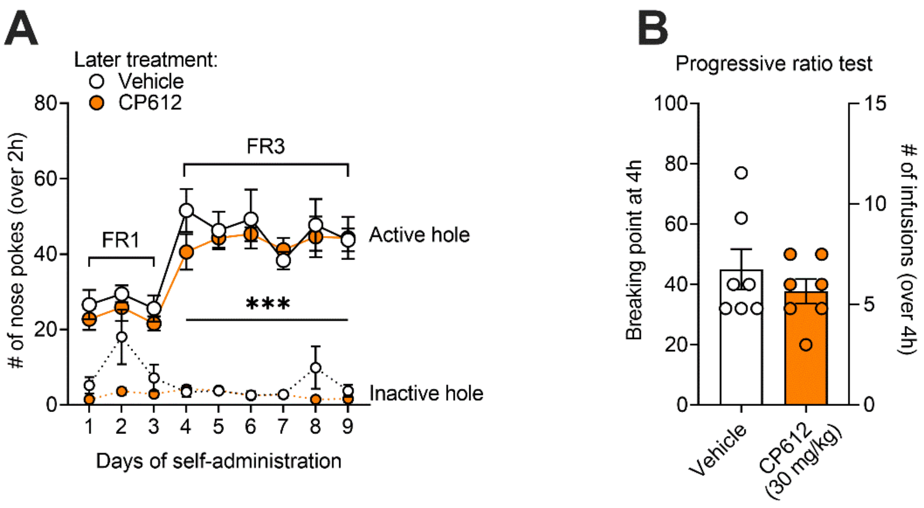


Figure S1. CP612 does not alter subsequent morphine self-administration. (A) Rats acquired morphine self-administration as attested by greater responding in the active hole vs. the inactive hole [$F_{\text{Hole type}}$ (1,12) = 264.68, $p < 0.001$] and this occurred to a similar extent in rats that would later receive vehicle or CP612 [$F_{\text{Hole type x PKC inhibition}}$ (1,12) = 0.012, $p > 0.91$]. The average intake of morphine was 13.4 infusions, and this intake was similar in rats that would later receive vehicle or CP612 [$F_{\text{PKC inhibition}}$ (1,12) = 0.182, $p > 0.79$; $F_{\text{Time x PKC inhibition}}$ (8, 96) = 0.47, $p > 0.88$]. (B) During the progressive ratio tests, the ratio to obtain an infusion was increased progressively within a self-administration session. Administration of CP612 (30 mg/kg, i.p.) 18h prior did not modify self-administration. This was measured as breaking point (highest ratio reached to earn an infusion of drug) and number of self-infusions $F_{\text{PKC inhibition}}$ (1,12) = 0.87, $p > 0.36$ and = 0.80, $p > 0.39$ for breaking point and infusions, respectively rats (n = 7 per group). Data are mean and SEM, dots are scores from individual rats. ***$p < 0.001$ compared with the inactive hole and with responding in the active hole during FR1.

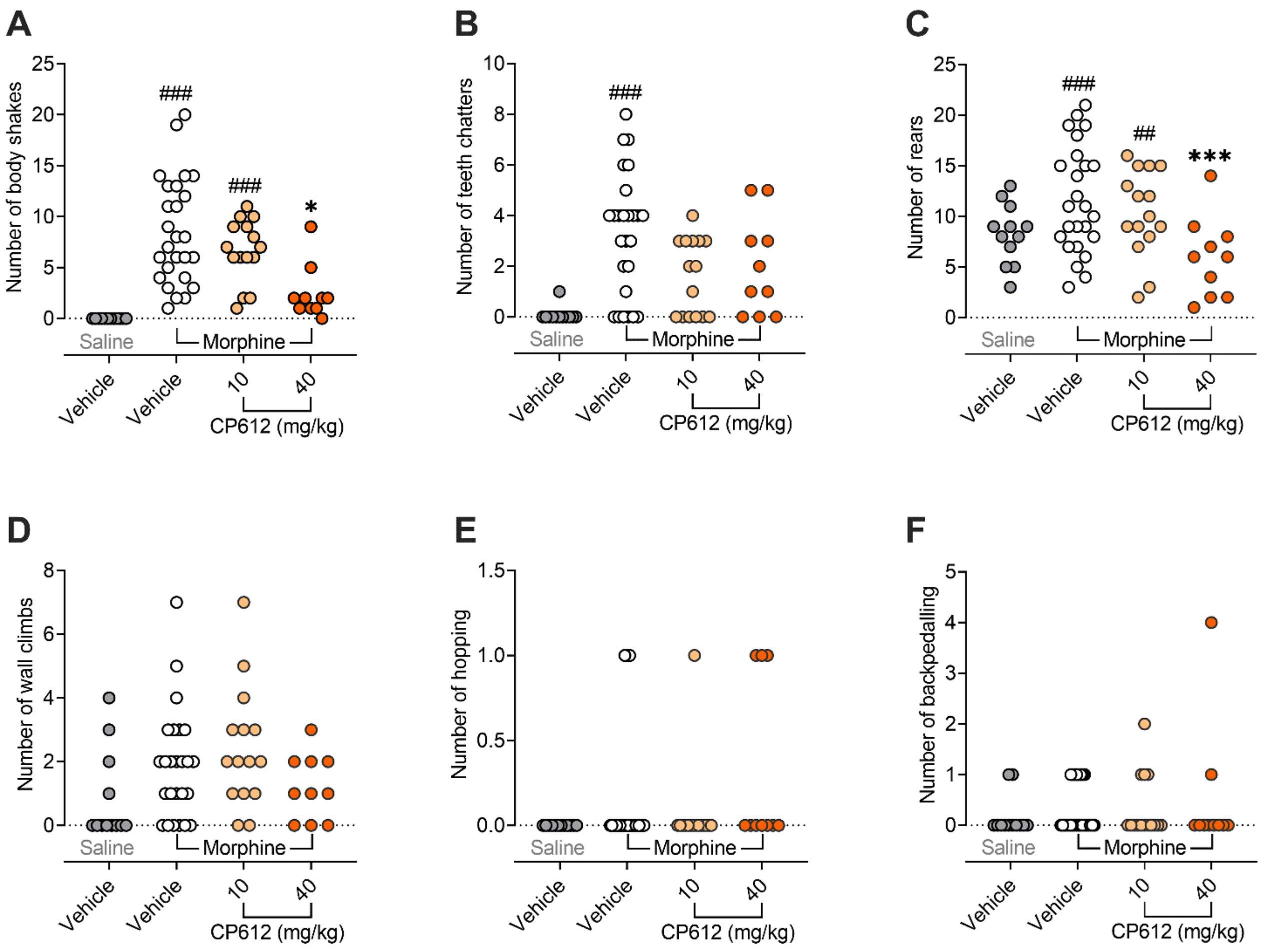


Figure S2. Morphine withdrawal precipitated by naloxone. Male mice received repeated injections of saline (10 mL/kg, i.p.) or morphine (20-100 mg/kg, i.p.) twice daily for 5 days. On day 6, during the conditioning day, they were pretreated with vehicle control (10 mL/kg, i.p.) or CP612 (10 mg/kg or 40 mg/kg, i.p.) followed by an injection of morphine (100 mg/kg, i.p.) 4h later. After 2h, they received an injection of naloxone (5 mg/kg, i.p.) to precipitate withdrawal. Withdrawal scores were analyzed with Kruskal Wallis test, due to the non-parametric nature of the data. Compared with mice that received repeated injections of saline, mice that received repeated injections of morphine showed withdrawal signs precipitated by naloxone for (A) body shakes [H (3, n = 63) = 37.216, $p < 0.001$], (B) teeth chatter [H (3, n = 63) = 37.16, $p < 0.001$], (C) rearing [H (3, n = 62) = 11.49, $p < 0.01$], and (D) wall climbing [H (3, n = 62) = 7.84, $p < 0.05$], but not (E) hopping [H (3, n = 63) = 4.79, $p = 0.188$], or (F) backpedaling [H (3, n = 63) = 0.81, $p = 0.848$]. Only the high dose of CP612 (40 mg/kg, i.p.) attenuated some of these withdrawal signs (##$p < 0.01$ and ###$p < 0.001$ compared with saline + vehicle; *$p < 0.05$ and ***$p < 0.001$ compared with morphine + vehicle). Dots are scores from individual mice.
